# A New Genus Of Microsporidian Parasite (Hepatosporidae; Micro-Sporidia) Found In The Oocytes Of Ribbon Worms From The North Pacific Genus Maculaura (Heteronemertea; Nemertea)

**DOI:** 10.1101/2021.04.04.438387

**Authors:** Kara M. Robbins, Svetlana A. Maslakova, George von Dassow

## Abstract

An intracellular microsporidian parasite was first observed within oocytes of *Maculaura alaskensis*, a small pilidiophoran nemertean, commonly found on sandflats along the Pacific coast of North America. Infected oocytes have large vesicles containing dozens to hundreds of diplokaryotic, ellipsoid spores measuring 1.3 by 2.3 μm. A partial small subunit nuclear ribosomal 18S gene sequence isolated from the microsporidian does not match any known microsporidian sequences in the public databases. Phylogenetic analysis groups it with *Hepatospora eriocheir* in a sister clade to the Enterocytozoonidae. All the known life stages of this parasite are contained within a membranous envelope. This microsporidian was identified in *M. alaskensis, Maculaura aquilonia, Maculaura oregonensis*, and *Maculaura cerebrosa* in Coos Bay, Oregon, in *M. alaskensis* from Newport, Oregon, and in *M. aquilonia* collected in Juneau, Alaska. This is, to our knowledge, the first species of microsporidia found to directly infect nemertean host cells.

**Graphical abstract:** 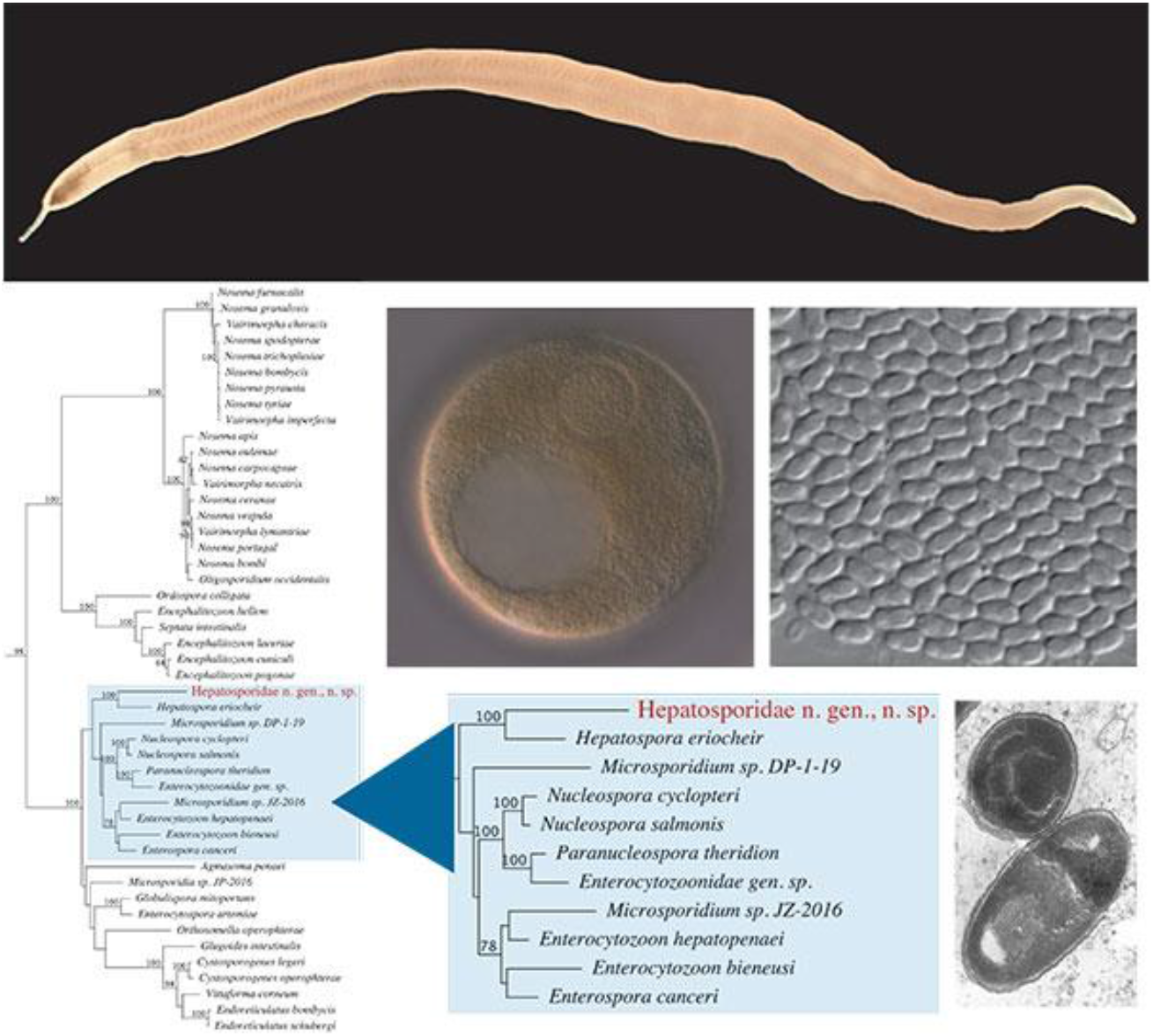

## 1. Introduction

Microsporidia are spore-forming intracellular parasites, considered either as a sister clade to or a basally branching group of fungi (Corradi and Keeling, 2009; Keeling and Fast, 2002). These specialized parasites have been found infecting protists and the cells of almost every animal phylum (Cali and Takvorian, 2014). Nearly 200 genera and 1,400 species have been described. Among the described genera, roughly 50 infect aquatic arthropods, and 21 infect other marine invertebrates and protists (Stentiford et al., 2013). These numbers certainly underestimate the diversity of microsporidia as some species infect multiple host species, and some hosts are susceptible to more than one microsporidian species. Microsporidian biologists hypothesize there is at least one microsporidian parasite for each species of animal on earth (Keeling and Fast, 2002).

Ribosomal gene sequences have been used for medical diagnosis and to establish modern microsporidian taxonomies (Vossbrinck et al., 2014; Vossbrinck and Debrunner-Vossbrinck, 2005). There are over 4,000 partial or complete small subunit ribosomal RNA 18S (SSU) reference sequences available for microsporidia in GenBank. Phylogenetic analyses of the SSU genes reveal five distinct clades (Vossbrinck and Debrunner-Vossbrinck, 2005). Clade IV is composed almost entirely of parasites of aquatic crustaceans and fish, with one notable exception: *Enterocytozoon bieneusi* causes microsporidiosis in immunocompromised human patients and is thought to be zoonotic (Stentiford et al., 2011). Within Clade IV, the family Hepatosporidae has been proposed following the discovery of *Hepatospora eriocheir*, a microsporidian infecting Chinese mitten crabs (*Eriocheir sinensis*) in Europe (Stentiford et al., 2011; Wang and Chen, 2007). At present, *H. eriocheir* is the only described member of this proposed family (Bojko et al., 2017; Stentiford et al., 2011).

Here we describe a new microsporidian infecting oocytes of ribbon worms in the genus *Maculaura* along the Pacific coast of North America. *Maculaura* live exclusively in marine and estuarine environments and are found under rocks or in sand within tidal mudflats (Hiebert and Maslakova, 2015). These worms range from Alaska to Southern California and have also been found in the Sea of Okhotsk in Russia (Coe, 1905; Hiebert and Maslakova, 2015). We find, with some frequency, oocytes of *M. alaskensis* or *M. aquilonia* containing large vesicles filled with small refractile ovoid objects that have diplokaryotic nuclei. We have determined these vesicles contain spores from a previously undescribed microsporidian that belongs to the family Hepatosporidae based on molecular phylogenetic and morphological analysis.

## 2. Materials and Methods

### 2.1 Collection

Live adult *Maculaura* and other nemerteans were collected from sandy mudflats using shovel and hand between July 2016 and October 2018 (ODFW collecting permits 202453, 21204, and 21962). The specimens came from mudflats we refer to as Portside (43°20’33 N, 124°19’27” W), Fisherman’s Grotto (43°20’32” N, 124°19’5” W), and Metcalf Marsh (43°20’10” N, 124°19’33 W) in Charleston, Oregon and Hatfield Marsh (44°37’4’ N, 124° 2’40” W) in Newport, Oregon. In the laboratory, individual *Maculaura* were separated into finger bowls filled with sea water filtered to 0.2 μm (FSW). All bowls were kept immersed in flowing seawater tables at ambient water temperature 13-18° C. Each individual *Maculaura* was given a unique designation and used for the following experiments. Adult *Maculaura aquilonia* tissue samples, collected from Juneau, Alaska by Terra Hiebert in August of 2014, preserved in 95% ethanol, and stored at −20° C, were also included in this study (Hiebert and Maslakova, 2015). Experiments were performed at the Oregon Institute of Marine Biology unless otherwise indicated.

### 2.2 Light Microscopy

Ripe female *M. alaskensis, M. aquilonia, M. magna*, and *M. cerebrosa* were placed in a petri dish with FSW. Ripe female *M. oregonensis* were not available for observation. A ~1 cm fragment was removed from the posterior end and oocytes were dissected from ovaries using a razor blade and fine forceps. Oocytes were examined with an Olympus BX51 DIC microscope for overt signs of infection.

### 2.3 DNA Extraction, PCR, and Sequencing

Visibly infected oocytes were collected in sets of 100 and frozen at −80° C for subsequent PCR testing. Samples were extracted using the Qiagen QIAamp Biostic Bacteremia DNA Kit with the following adjustments. A 0.5 mm PowerBead Tube was used for the first incubation and subsequent vortexing step. Oocytes were incubated in Buffer MBL (Qiagen) at 70° C for 20 minutes mixing at 1,400 rpm.

Universal microsporidian primers for the small subunit ribosomal RNA (SSU) and RNA Polymerase II (RPB1) genes were used to amplify a 1280 base pair region of the SSU gene and a 885 base pair region of RPB1 from the unknown microsporidia (Table 1). All PCR reactions were performed with Promega GoTaq DNA polymerase using a 20 μL reaction volume and the following variable parameters: 0.5μM or 1 μM of each primer, and 2 μL of template DNA. Reaction conditions consisted of: initial denaturation for 2 minutes at 95°C followed by 34 cycles of 40 seconds at 95°C, 40 seconds at 43° C for SSU or 56° C for RPB1, and 1 minute 15 seconds for SSU or 1 minute 5 seconds for RPB1 with a final elongation step of 2 minutes at 72°C.

**Table 1.**
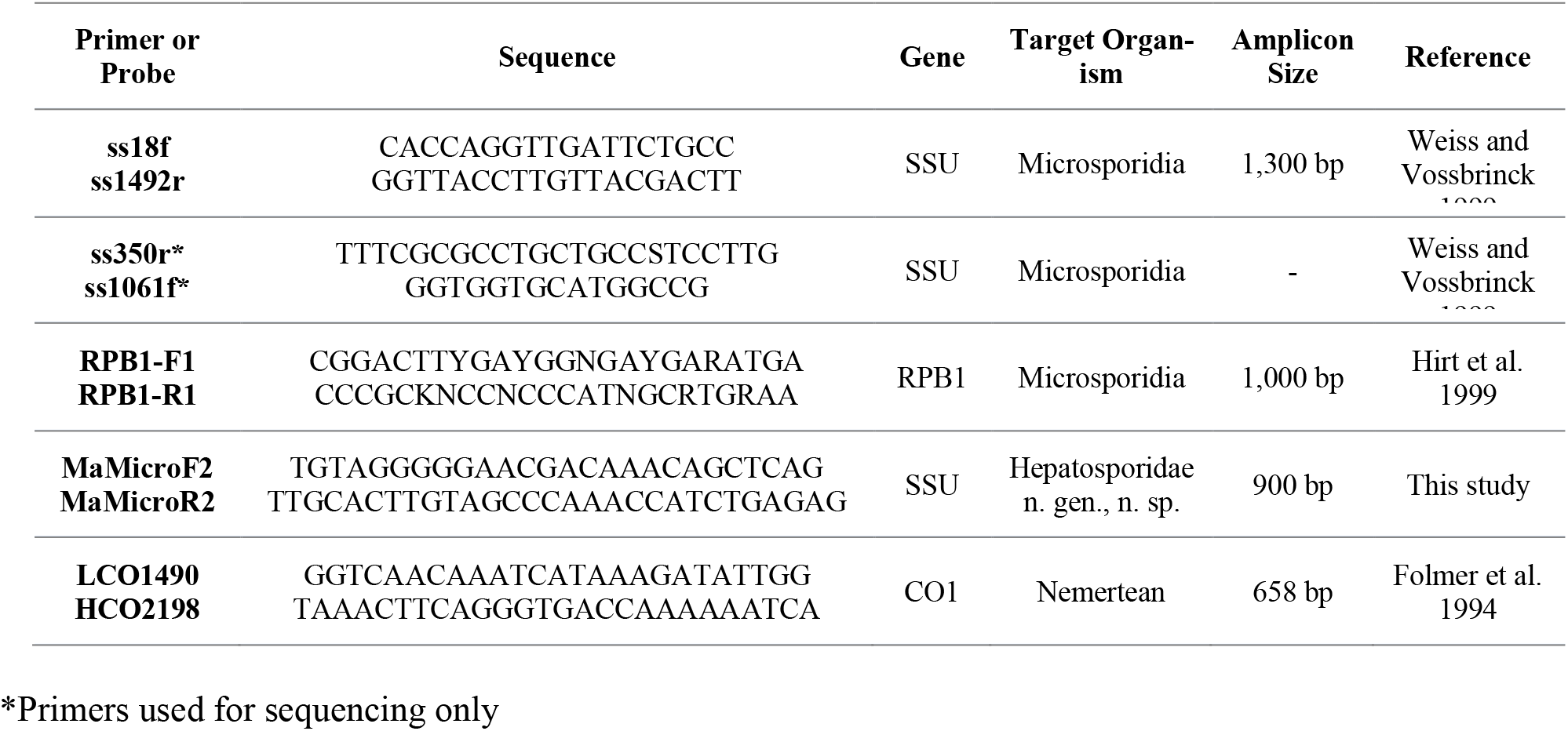
List of all primers used for PCR and sequencing. All primers listed read 5’ to 3’.

The amplified products were ligated into Promega pGEMT+ Vector Systems and cloned using Invitrogen DH5a cells. Colonies were selected and re-amplified with the same universal primers mentioned above, purified, and sent for Sanger Sequencing. All sequencing was performed by Sequetech Corporation (Mountainview, CA) using the same universal primers along with ss350r and ss1061f for the SSU gene. Sequences were trimmed and aligned using Geneious software version 11.0.4. Consensus sequences were run through the BLASTn and BLASTx databases.

### 2.4 SSU Phylogenetic Analysis

For phylogenetic analysis, an additional 137 microsporidian SSU sequences were downloaded from the NCBI database with emphasis on species from Clade IV (Vossbrinck and Debrunner-Vossbrinck, 2005). These sequences, plus the consensus SSU sequence for the unknown microsporidian were aligned with MAFFT as implemented in Geneious using default parameters. J-Model test was used to determine the best fit model for maximum likelihood (ML) analysis. A ML tree was created using Phyml with a GTR+I+G Model using 200 bootstrap replicates implemented in Geneious. The same alignment and model were used to create a Bayesian inference phylogeny (using Mr. Bayes plug in Geneious).

### 2.5 Transmission Electron Microscopy (TEM)

Sets of oocytes including some number of visibly infected individuals were fixed in 2% glutaraldehyde in 0.2 M sodium cacodylate buffer (SCB, Electron Microscopy Sciences) for one hour and post-fixed in 2% osmium tetroxide in SCB for one hour (Bozzola and Russell, 1999). Oocytes were infused and embedded in LR white resin with LR accelerant at the Center for Advanced Materials Characterization (CAMCOR) at the University of Oregon. Ultra-thin sections, approximately 90-100 nm thick, were prepared with freshly broken glass knives on an LKB Ultramicrotome III and mounted on uncoated 300 mesh copper grids. Sections were stained with 4% uranyl acetate and Reynold’s lead citrate (Reynolds, 1963) then imaged with the FEI Tecnai G2 Spirit TEM.

### 2.6 Histology

Midbody regions from adult female *Maculaura* with visibly infected oocytes, females with visibly uninfected oocytes, and adult male *Maculaura* were fixed and stained for histological examination. Specimens were relaxed in a 1:1 mixture of 0.33 M MgCl_2_ and FSW at 4°C (30 minutes), fixed with 10% formalin in phosphate buffered saline (PBS) (24 hours), post-fixed in Hollande’s Bouin fixative (72 hours), dehydrated through a graded ethanol series (30%, 50%, and 70% for 10 minutes each), then rinsed daily with 70% ethanol until the rinsing alcohol remained colorless. Tissues were stored in 70% ethanol for further processing.

Samples were dehydrated, cleared, and paraffinized (Richard-Allan Scientific Paraffin 9, 56°C melting point) in an automated tissue processor at the University of Oregon Histology Laboratory. Once embedded in paraffin, tissues were sectioned at 7 μm then mounted on glass slides. Slides were deparaffinized in xylene (2, 3 minute rinses), rehydrated with a graded ethanol series (100% twice, 95%, 85%, 70%, and RO water twice at three minutes for each step), and stained using Weber’s Chromotrope (6 g chromotrope 2R, 0.15 g fast green, and 0.7 g phosphotungstic acid in 3 mL glacial acetic acid) (Weber et al., 1992; Ghosh et al., 2014). Stained slides were dehydrated using a reverse of the ethanol series described above, cleared in xylene (2, 3 minute rinses), mounted in Permount mounting medium, and examined with an Olympus BX51 DIC microscope. Histology images were taken using a SPOT Insight 12 MP camera and SPOT Basic 5.6 software.

### 2.7 Larval Development Observations

*M. alaskensis* larvae from females carrying some fraction of infected oocytes were grown in 150 mL finger bowls filled with FSW, and set in running sea water tables at 13-18° C. Water was exchanged every three days and larvae were fed cultured *Rhodomonas lens* after each water change. At 7, 14, 30, and 40 days of development individuals were photographed and frozen at - 80° C for PCR testing. Additionally, following metamorphosis (between 36 and 41 days), juveniles were collected, frozen, and PCR-tested for infection.

### 2.8 Diagnostic PCR for Detection of the New Microsporidian

A specific primer set was designed to amplify a 900-base pair fragment of the unknown microsporidian SSU sequence (Table 1). Live adult *Maculaura* collected between January and October of 2018 were examined for their reproductive status (not ripe, ripe female, ripe male), then a 2-3 mm^3^ cross-section was removed from the posterior 1/3 of the animal and frozen at - 80° C. Between animals, tools and petri dishes were washed with 10% bleach, rinsed in RO water, UV treated for 20 minutes on each side, then washed and rinsed in a separate 10% bleach and RO container to prevent cross contamination. Larvae and newly metamorphosed juveniles from developmental studies along with the preserved adult specimens from Juneau, Alaska in 2014 were also tested.

Adult tissues frozen in FSW were removed from the freezer and rinsed once in 1 mL nuclease-free water prior to DNA extraction. Samples stored in ethanol were rinsed three times in 1 mL nuclease-free water over 10 minutes before extraction. Following rinsing, adult tissues were ground with a plastic pestle in a microcentrifuge tube. All samples were extracted using the QIAamp Biostic Bacteremia DNA Kit following the protocol described previously. Larvae and new juveniles were extracted using 1/4 volume of glass beads and 1/3 volume of Buffer MBL, IRS, and BB. All other volumes remained the same. Amplification was carried out with GoTaq DNA Polymerase (1U per reaction) with 0.5μM of each primer and 2 - 8 μL of template DNA. Cycling conditions consisted of an initial denaturation at 95° C for 2 minutes followed by 34 cycles of 40 seconds at 95° C, 40 seconds at 52° - 56° C, and 55 seconds at 72° C followed by 2 minutes at 72°C. Results were assessed by gel electrophoresis. Of the samples which produced a band of the expected size, 40% were purified and sent for Sanger sequencing with the same specific primers. Sequences were trimmed, assembled, and compared to the original partial SSU sequence using Geneious.

### 2.9 DNA Barcoding of Infected Nemerteans

The species identity of all samples which tested positive for microsporidian infection by PCR was confirmed by DNA barcoding using universal metazoan Cytochrome Oxidase I primers (Folmer et al., 1994), because species of *Maculaura* are cryptic (difficult to tell apart morphologically). Reactions were run as above with the following exceptions: 2 μL of template DNA, annealing temperature of 45° C, and elongation time of 1 minute. Samples were purified and sequenced in both directions by Sequetech Corporation. The resulting sequences were trimmed and assembled in Geneious and analyzed using BLASTn. All samples and reference sequences were aligned using MAFFT as implemented in Geneious, and a Neighbor-Joining tree was constructed including sequences of the five described species of *Maculaura* and an outgroup.

## 3. Results

### 3.1 Initial Observations

Spore-filled parasitophorous vesicles (PSVs) were discovered in oocytes of *M. alaskensis* (Figure 1A) and *M. aquilonia* but not found in *M. cerebrosa* or *M. magna* (*M. oregonensis* oocytes were not observed). These PSV’s are often found closely associated with meiotic or mitotic spindles of the developing oocyte and embryo (Figure 1A–1C) and they contain uniformly shaped ellipsoid spores (Figure 1A). Live staining with Hoechst 33342 (a DNA dye) revealed diplokaryotic nuclei contained in each individual spore (Figure 1B and 1C). Oocytes from four infected *M. alaskensis* females contained anywhere from one to twelve PSVs in the cytoplasm with an average of 2.36 PSVs per infected oocyte. Between 5 and 16% of oocytes from an infected female had visible spore-filled vesicles (Table 2). The size of the PSV and the number of spores contained within the vesicles varied dramatically: PSV diameters ranged from 6 to 30 μm, so volumes ranged from ~0.1-10 picoliters. Earlier life stages were also observed within PSVs, but at a much lower frequency than spore-filled vesicles.

**Figure 1:**
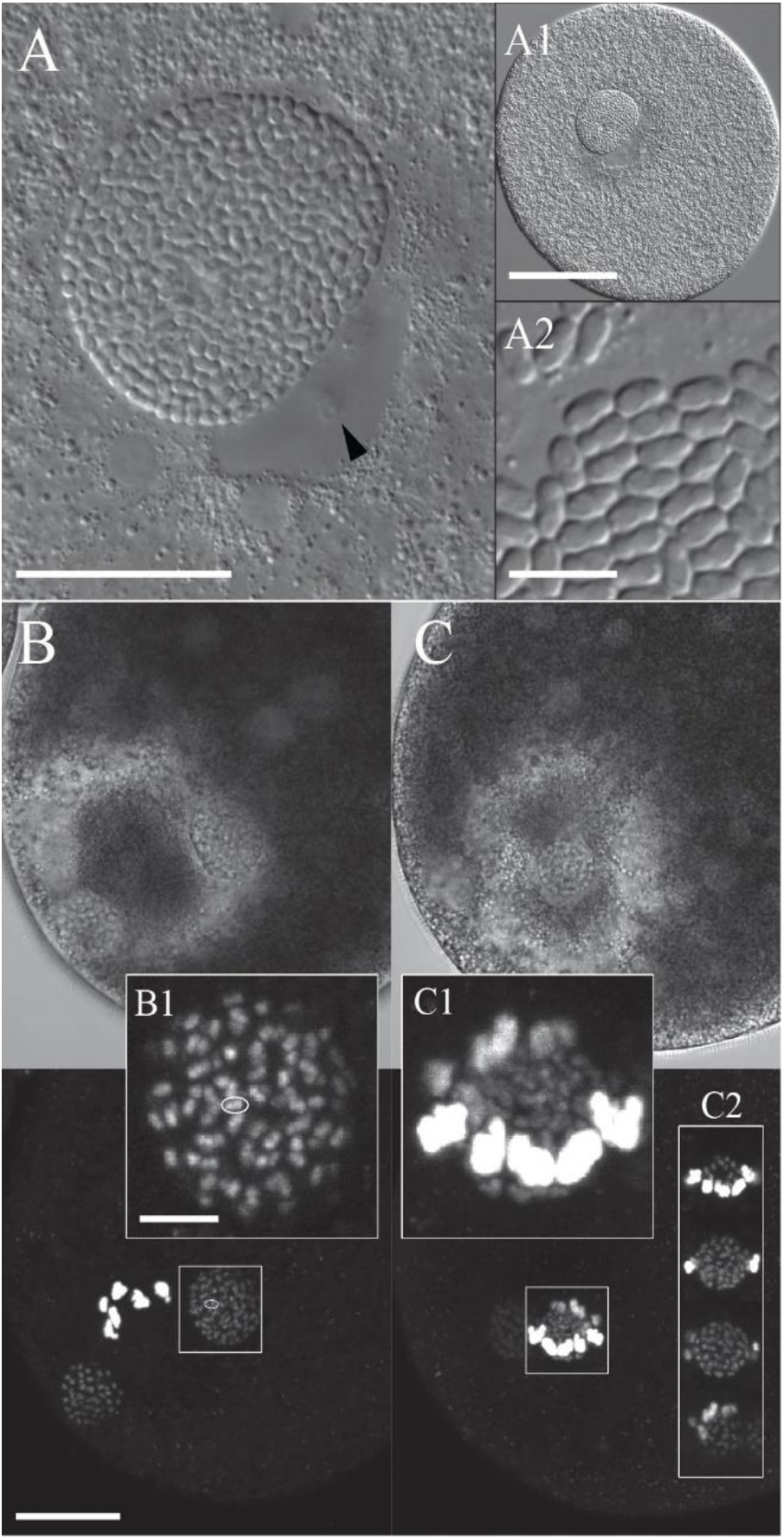
DIC and confocal images of initial observations of infected oocytes. A) High-magnification view of PSV associated with the meiotic spindle in a living, compressed oocyte. Arrowhead indicates host chromosomes. SB = 20 μm. A1) Low magnification image of infected oocyte (SB = 50 μm). A2) A 2x blown-up image of spilled spores after the oocyte was crushed under the coverslip (SB = 5 μm). B, C) Living oocytes stained with Hoechst demonstrate diplokaryotic spores in PSVs associated with the meiotic spindle (B) and an average-sized PSV embedded within the meiotic spindle (C). SB = 20 μm. Insets B1 and C1 are 3x blow-ups of the boxed region (projections of 10 and 22 consecutive 0.6-micron confocal sections, respectively. SB = 5 μm); inset C2 shows successive focal heights within the boxed region, to show that the PSV is surrounded by host bivalents.

**Table 2:** Brief summary of epidemiological observations from *Maculaura alaskensis* females collected in 2017. Includes a count of infected oocytes versus uninfected oocytes in each female as well as the PSV count per oocyte.

| Specimen ID | Infected Oocytes | Uninfected Oocytes | % Infected | Average PSV Count per Oocyte | Max PSV Count From Single Oocyte |
| --- | --- | --- | --- | --- | --- |
| MaPsIn 17-1 | 32 | 170 | 15.8% | 2.844 | 10 |
| MaPsIn 17-2 | 32 | 222 | 12.6% | 2.5625 | 12 |
| MaPsIn 17-3 | 10 | 182 | 5.2% | 1.8 | 8 |
| MaPsIn 17-5 | 13 | 88 | 12.9% | 2.2308 | 8 |

### 3.2 Sequences and phylogenetic analysis

Four, 1,280 bp small subunit ribosomal RNA (SSU) sequences (GenBank accession numbers MT302614 - MT302617) were amplified and used for identification. These sequence had between 74.68% and 76.27% sequence similarity to *Microsporidia* sp. DP-1-19 (NCBI Accession AF394528, 91-95% query coverage) grouping this organism within Clade IV and class Terresporidia (Vossbrinck and Debrunner-Vossbrinck, 2005). Five, 881 to 890 bp RPB1 sequences (GenBank accession numbers MT360642-MT360646) had a query coverage no higher than 15% when analyzed with BLASTn. Using BLASTx, the protein sequences matched *Hepatospora eriocheir* (between 97-98% query coverage and 54.95-56.9% identity), another member of Clade IV and within the proposed family Hepatosporidae (Bojko et al., 2017; Stentiford et al., 2011; Wang and Chen, 2007).

Maximum Likelihood and Bayesian inference (Figure 2) phylogenies of 138 microsporidian SSU rDNA sequences group the unknown microsporidian in Clade IV within the monotypic family Hepatosporidae (its sole member until now being *Hepatospora eriocheir*) with strong support (100% bootstrap and 1.0 posterior probability). The unknown microsporidian and *H. eriocheir* form a sister clade to the Enterocytozoonidae with a posterior probability of 1.0 and 70% bootstrap support (Figure 2, Supplemental Files 1 and 2). Based on a unique SSU rDNA sequence, and its position on the phylogenies, we propose a new genus and species *Oogranate pervascens* gen. nov., sp. nov. within the family Hepatosporidae.

**Figure 2:**
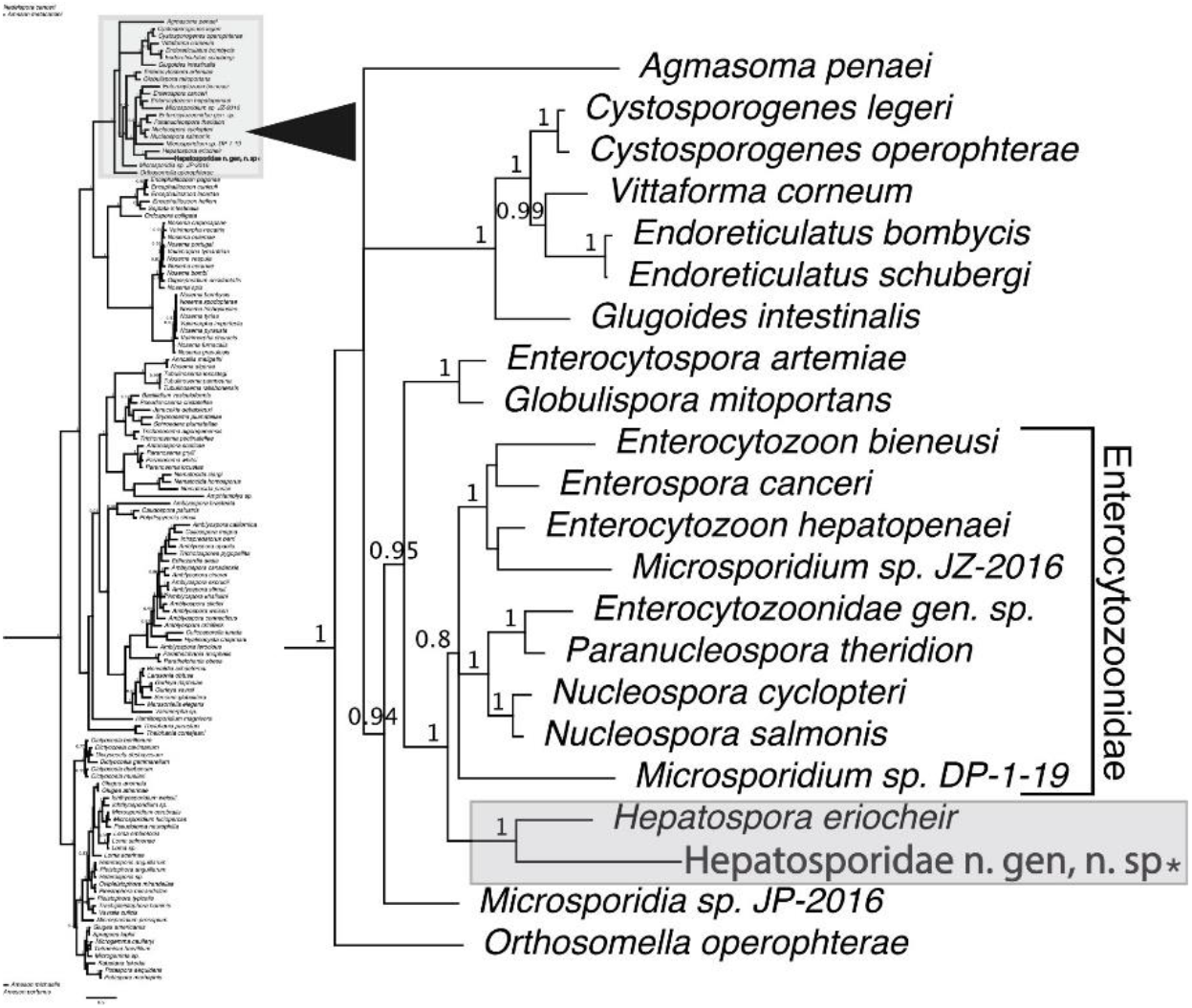
A clade from the Bayesian inference phylogeny of 138 microsporidian SSU sequences including the unknown microsporidian from this study (bold font, asterisk. The Hepatosporidae clade is highlighted in grey. Posterior probabilities less than 0.75 are not shown. See Supplemental File 1 for full Bayesian inference phylogeny.

### 3.3 Morphology

Analysis by TEM showed the spores are coated with an electron-dense exospore, an electron-lucent endospore, and a plasma membrane. Spores also contain a polar tube, two nuclei, and a posterior vacuole. The polar tube measures 80 nm in diameter and is arranged in a single layer of six to seven coils. Spores are 2.3 μm long and 1.3 μm wide on average (Figures 3A and 3B). Vesicles within the cytoplasm were observed either docking with or leaving from the PSV (Figure 3B). Complete spores were always found within a PSV, and earlier life stages were not observed with TEM. In one case, a single matured spore with an extended polar tube was found inside a PSV (Figure 3C).

**Figure 3.**
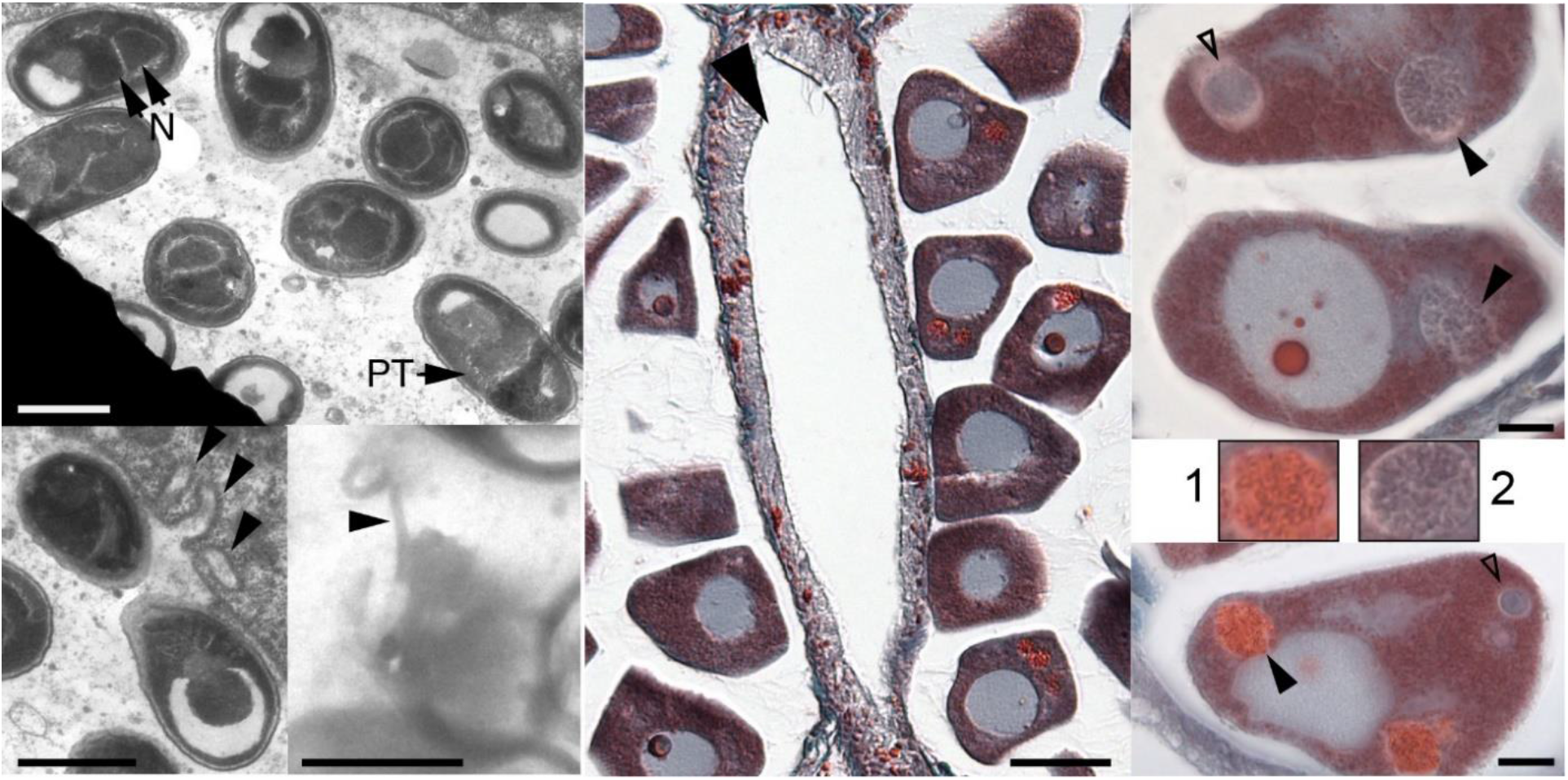
TEM and Histology images of infected oocytes containing PSV’s. A) TEM image of spores found in an infected oocyte. Nuclei (N) and polar tube (PT). B) Vesicles docking or leaving the PSV (arrowheads). C) Mature spore with extended polar tube (arrowhead). All TEM images SB = 1 μm. D) Low-magnification histology image of infected female *M. alaskensis tissue*, SB = 50 μm. Arrowhead indicates gut diverticulum lying between two separate ovaries. Ovary on the right contains several infected oocytes, ovary on the left contains none. E, F) Two oocytes with PSVs in various stages of microsporidian development, SB = 10 μm. E) Solid arrowheads 1 indicate potential sporoblasts (inset 1). Unfilled arrowhead points to an earlier developmental stage. F) Solid arrowhead points to a PSV full of complete spores (inset 2). Open arrowhead indicates an earlier developmental stage.

In histological sections, Weber’s Chromotrope stained the exospores bright red and demonstrated a visible, uniform internal structure within each complete spore (Figure 3F). Other life stages of the parasite were also detected and stained blue (Figure 3E and 3F). Individual oocytes could be found carrying multiple separate PSVs at different stages of microsporidian development (Figure 3D - 3F). Often, nearly all oocytes in a single ovary would be infected, while the next ovary would have no signs of infection (Figure 3D).

Nemertean muscles, epithelium, and other tissues of the adult *Maculaura* stained blue. Certain structures in the epidermis and dermis of *Maculaura* stained red (likely, gland cells), but were distinguished from microsporidian spores by their lack of distinctive internal features. Using the criteria described here, infections were only detected within PSVs of infected oocytes and never observed within adult tissues.

### 3.4 Diagnostic PCR and Identification of Nemertean Host Species

Between January and October of 2018, a total of 70 adult nemertean worms from the genus *Maculaura* collected in Charleston, Oregon were tested for microsporidia by PCR with novel microsporidian species-specific primers. Of these, 33 tested positive (produced a visible band of expected size as assessed by gel electrophoresis). Positive individuals included ripe females, ripe males, and individuals without ripe gametes. They included representatives from four of five species of the genus *Maculaura: M. alaskensis, M. aquilonia, M. oregonensis*, and *M. cerebrosa* (Figure 5). Three samples from the fifth species, *M. magna*, were negative for infection by PCR (data not shown). Of the Charleston *Maculaura*, 41.4% tested positive for infection by PCR (43.3% excluding *M. magna*).

**Figure 4.**
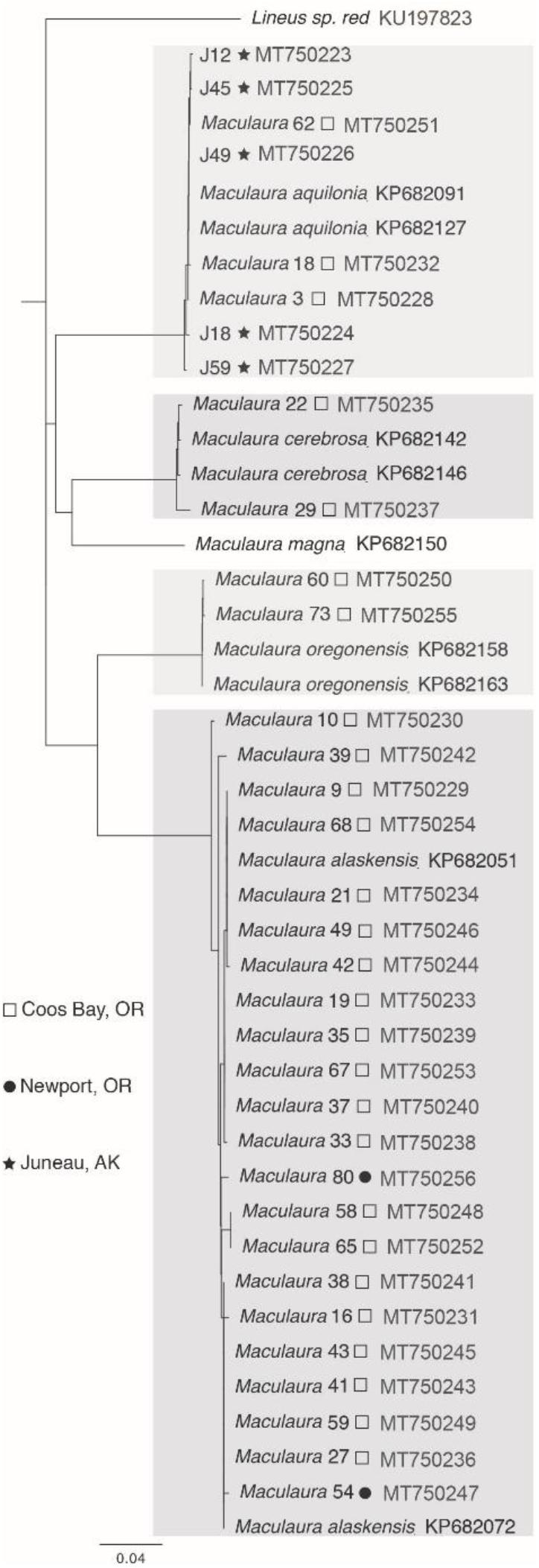
Neighbor-joining tree of *Maculaura* host COI sequences used to identify nemertean hosts positive for infection with *Oogranate pervascens* gen. nov. sp. nov, along with reference sequences from GenBank. Shaded boxes correspond to different species of *Maculaura* from top to bottom as follows: *M. aquilonia, M. cerebrosa, M. oregonensis*, and *M. alaskensis*.

**Figure 5:**
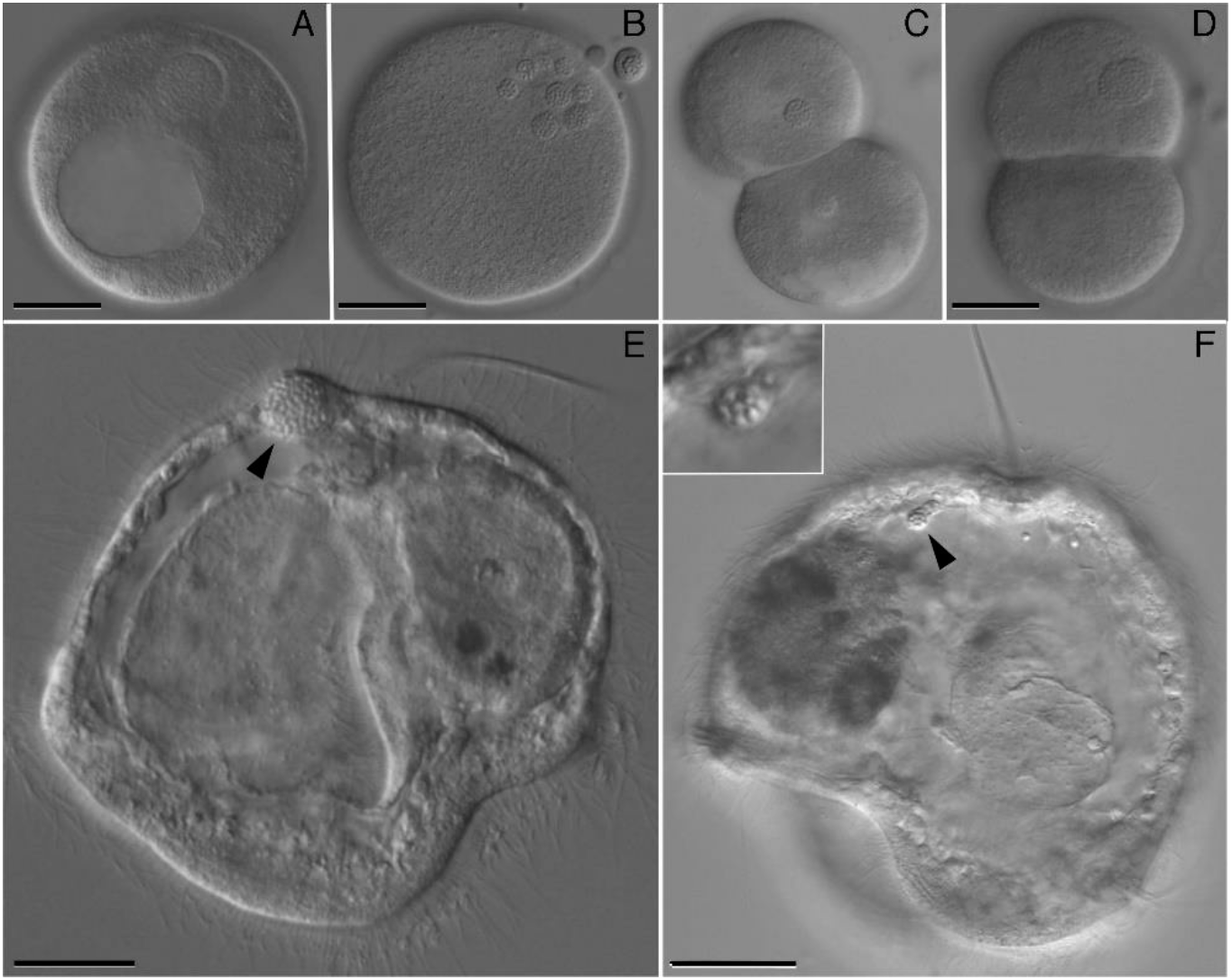
Observations of *Maculaura alaskensis* embryos and larvae developing from infected oocytes. A) Oocyte with one large PSV filled with spores outside of the germinal vesicle. B) Mature egg with several PSV’s including one PSV inside the first polar body. Egg compressed tightly with coverslip to show extent of infection. C) Infected egg immediately following first cleavage. D) Infected egg containing 2 large PSV’s (one out of focus) following first cleavage. E) A 3-day old pilidium carrying a large spore-filled mass (solid arrowhead). F) A 2-week-old pilidium with an infected mesenchyme cell (solid arrow-head and inset). SB = 25 um.

Of the 17 *Maculaura* specimens collected from Newport, Oregon, two *M. alaskensis* tested positive. All the worms collected from Alaska in 2014 were previously identified as *M. aquilonia* by Terra Hiebert (Hiebert and Maslakova, 2015). Of the 25 samples of Alaskan *M. aquilonia* tested, five were positive for infection. The Neighbor-Joining tree of nemertean CO1 sequences (Figure 4) confirms results obtained from BLASTn. The microsporidian novel primer PCR test were verified by Sanger sequencing. Thirteen of the 34 (38.2%) samples which produced a band of expected size were sent for sequencing. All 13 sequences matched the SSU ribosomal sequence with 99-100% similarity, regardless of host species or sampling location. In summary, we positively detected the same novel microsporidian in males, females, and unripe individuals from four out of five *Maculaura* species in three distinct locations; limited sampling has so far failed to detect the presence of this microsporidian in other co-occurring nemertean species including: *Cerebratulus* spp., *Paranemertes* spp., *Emplectonema* spp., and *Zygonemertes* spp.

### 3.5 Development of infected eggs and larvae

Infection does not entirely preclude apparently normal larval development. Some fraction of infected oocytes, even those containing numerous or large PSVs, were capable of maturing (Figure 5B), fertilizing, initiating development (Figure 5C and 5D), and growing into swimming and feeding pilidia (Figure 5E and 5F). Other heavily infected oocytes often failed to mature, fertilize, or begin cleaving.

A visibly infected pilidium with a large cyst containing spores was found after two weeks of development. This 2-week-old pilidium was tested to verify the use of diagnostic PCR on infected larvae. This visible infection, carried inside a mesenchyme cell of the pilidium, was confirmed positive by PCR (Figure 5F). To check for cryptic infection, pilidia lacking visible spores were tested by PCR. None of 12 pilidia at various stages of development, nor any of seven metamorphosed juveniles, tested positive for microsporidia infection by PCR.

## 4. Systematics

### 4.1 Oogranate pervascens *gen. nov., sp. nov*

#### Description

Refractile ellipsoid spores are found densely packed into PSVs (Figure 1A). Spores measure 2.3 X 1.3 μm (2μm^3^) in TEM prepared oocytes. The spores are diplokaryotic (Figure 1B inset, Figure 1C, and Figure 3A) and contain 6-7 polar tube coils in a single row (figure 3A). The spore coat has an electron-dense exospore and a thin electron-lucent endospore. All life stages are housed within a vesicle capable of interacting with, but separate from, the host-cell (Figure 3B).

#### Diagnosis

Vesicles containing all life-stages are visible within oocytes of *M. alaskensis* and *M. aquilonia* by light microscopy (Figure 1A and Figure 5A). Spores are also visible by light microscopy inside mesenchyme cells of developing *M. alaskensis* pilidia (Figure 5F). Hoechst and propidium iodide (PPI) staining can be used for detecting microsporidian diplokarya in cytoplasm of the oocytes even inside whole mount adult tissues cleared with xylene. Multiple life-stages are visible in histological sections of ripe female *M. alaskensis*. Spores are also visible by TEM of prepared oocytes. Species-specific primers MaMicroF2 and MaMicroR2 (Table 1) can be used for nucleic acid-based diagnosis. Universal microsporidia primers ss18sf and 1492r as well as RPB1-F1 and RPB1-R1 can be used for comparison to full-length sequences under the following GenBank accession numbers: MT302614 to MT302617 and MT360642 to MT360646 (See Table 1 for all primer sequences and references).

#### Type host

*Maculaura alaskensis*.

#### Type locality

Coos Bay Estuary in Charleston, Oregon, U.S.A.

#### Site of infection

Cytoplasm of *M. alaskensis* and *M. aquilonia* oocytes, usually clustering close to the nucleus. Also, the cytoplasm of mesenchyme cells of developing *M. alaskensis* pilidia. Other sites of infection are unknown. Sites of infection within *M. cerebrosa* and *M. oregonensis* are unknown.

#### Etymology

We suggest the name *Oogranate pervascens*. The genus is named for the characteristic spore-filled PSVs within host oocytes (from Latin “granum” = seed and Greek “oon”= egg). Species name for the abundance infected oocytes found in infected females towards the end of their reproductive season, the number of spores produced, and in reference to the currently mysterious route of infection (from latin “pervadere” = permeate).

## 5. Discussion

### 5.1 Host Species

Of the five described *Maculaura* species, the only one where infection was not detected by any technique is *M. magna*, a worm aptly named for its large body size, compared to its congeners: the other *Maculaura* species average between 3 and 10 cm in length and 1-3 mm in width, whereas specimens of *M. magna* measure on average 20 cm long and 3-4 mm wide (Hiebert and Maslakova, 2015). Only three *M. magna* from Charleston, Oregon were tested by PCR, but if further sampling confirms that this congeneric nemertean does not host the newly-described microsporidian, body size suggests a biological rationale. Though this nemertean occupies a similar habitat to other *Maculaura* and has the same life history strategy (i.e., a maximally-indirect development within a pelagic larva), it may specialize on different prey species compared to its smaller relatives. If microsporidian infection in the other *Maculaura* is acquired from prey, this might explain why *M. magna* does not share this parasite with its congeners.

### 5.2 Parasite Load

Based on size and numbers of PSVs within infected *M. alaskensis* oocytes, parasite loads varied widely (Table 2). Spore numbers within oocyte PSVs ranged from 10^2^ to 10^4^. The average oocyte infection recorded from Coos Bay was 2.4 PSVs per oocyte and the median size was 1 picoliter (based on a PSV diameter of 12 μm). If an *M. alaskensis* oocyte has an approximate volume of about one quarter of a nanoliter (75 microns in diameter), then a moderate infection of 2.4 PSVs measuring 1 picoliter each might take up only 1% of the oocyte volume but produce 1.2X10^3^ spores in a single oocyte. The ability of this parasite to produce relatively high spore loads within a single PSV is likely due to infection within oocytes, which are one of the largest and best-provisioned cell types.

*M. alaskensis* infections are most apparent and contain heavy parasite loads. However, this finding may be due to higher sampling of this species. *M. cerebrosa* is another often-encountered species which tested positive for this microsporidian by PCR. However, we have not observed overtly infected oocytes in this host, which may indicate that infection load is lower or present in a different tissue type in this species. A fluorescent in-situ hybridization method or a more thorough histological sampling will be necessary to compare pathology between different host species.

### 5.3 Parasite Development

All the stages we have detected by histology in *M. alaskensis* – the proliferative stage (merogony), the spore-forming stage (sporogony), and the infective stage - are separated from contact with oocyte cytoplasm by an interfacial membrane (Figure 3A, 3B, 3E, and 3F). This membrane is likely of host origin since it is found throughout the life cycle (Weiss and Becnel, 2014).

Figure 6 depicts a hypothetical lifecycle for *Oogranate pervascens* based on observations of Weber’s Chromotrope stained histological sections of infected *M. alaskensis* females. Presumably, infection begins with injection of spore contents into the oocyte cytoplasm during oogenesis as so far we have not been able to determine the initial source of infection. The meront appears as a dense, uniformly blue stained, circular object within the PSV. The lack of visible boundaries within this object suggests merogony proceeds by plasmotomy, production of a single large cell with many nuclei. Once individual cells are separated from the larger plasmodium, they transition to spore formation (Cali and Takvorian, 2014; Vávra and Larsson, 2014).

**Figure 6.**
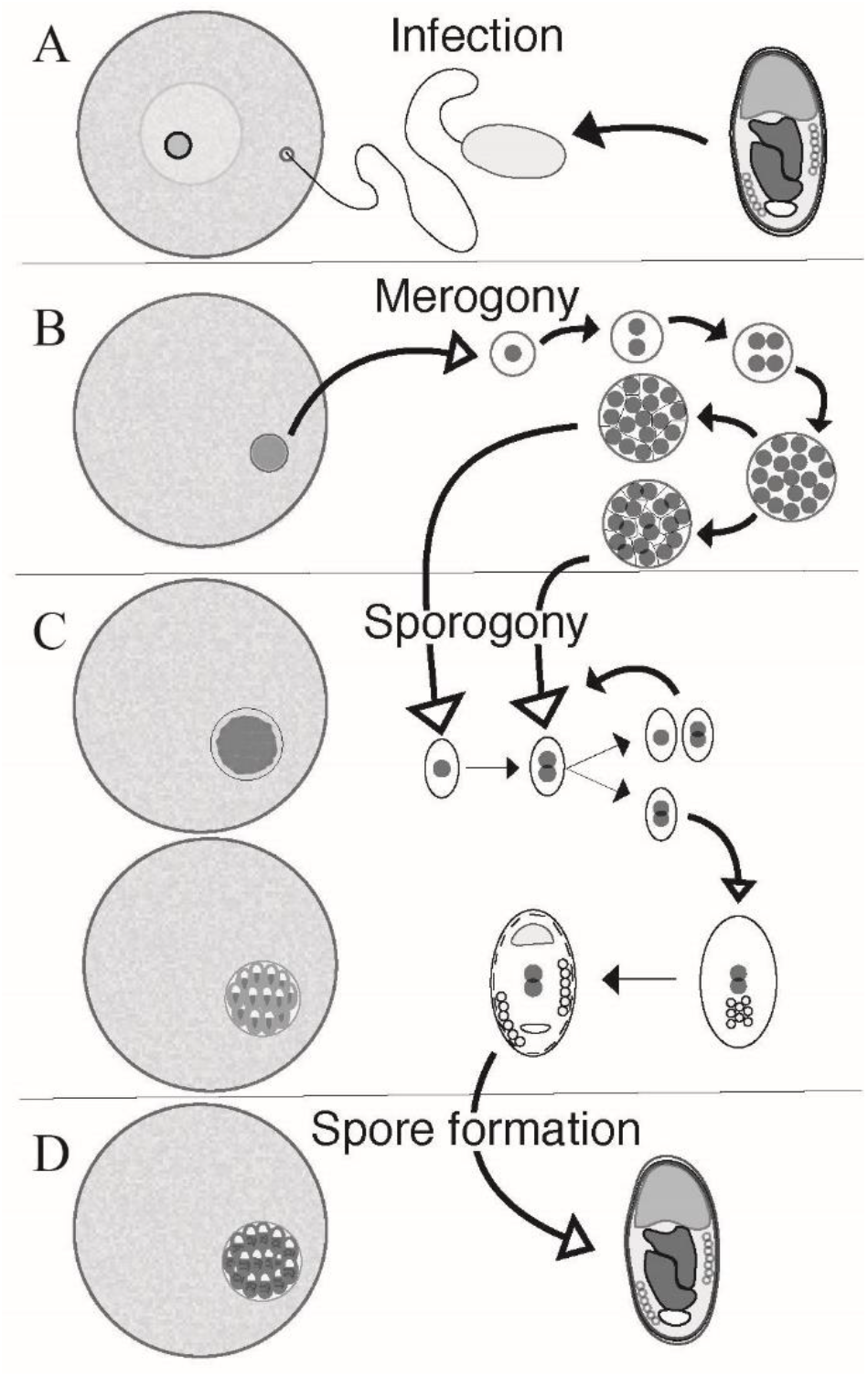
Hypothesized life cycle of *Oogranate pervascens*. Oocytes (left) show a histological representation of the lifecycle stage (right). A) Initial infection by injection of spore materials into oocyte cytoplasm and formation of PSV. B) Proliferation proceeds by plasmotomy, where many nuclei are produced inside a single cell body. The plasmodium is carved into individual unikaryon or diplokaryon cells marking the end of merogony. C) Sporogony initiates with division of newly formed cells. The daughter cells continue dividing, then eventually differentiate and form specialized spore contents. D) Spore formation is completed after secretion of the spore coat.

During sporogony, as in many other microsporidia, sporonts divide producing many spores. It appears the PSV expands outward during this stage, evidenced by a clear ring visible between the developing microsporidia and surrounding envelope, to give room for parasite growth (Figure 3E). Cells eventually stop dividing and begin to differentiate by developing the polar tube and spore coat, marking the end of sporogony. Sporoblasts appear as individual blue cells with uniform internal structure similar to the configuration observed in completed spores. Spore development concludes with the formation of a completed electron-dense coat which stains bright red with Weber’s Chromotrope stain. This inferred life cycle requires confirmation by TEM, and may proceed differently in other host cells or in the oocytes of other *Maculaura* species.

### 5.4 *Mode of Transmission in* Maculaura alaskensis

Mode of transmission, whether horizontal or vertical, is a key descriptor of microsporidian species. We have no firm evidence supporting vertical or horizontal transmission of this microsporidian. Using TEM, a single matured spore with extruded polar tube was observed (Figure 3C). In histological sections, several infected oocytes are usually found in the same ovary. This could mean that some fraction of spores mature within oocytes, resulting in parasite transmission between adjacent oocytes. Such a strategy is used by other microsporidian species to pass infection into neighboring cells (Cali and Takvorian, 2014). Oocyte-to-oocyte transmission seems plausible, but whether initial infection of an individual occurs by ingestion or inheritance is yet to be determined.

#### 5.4.1 Vertical Transmission

The fact that viable gametes carry spores suggests the possibility of vertical transmission. We can occasionally detect infection in developing larvae, but presence does not guarantee transmission to the juvenile. Our preliminary observations suggest that infected pilidia either fail to develop to metamorphosis or clear the infection before then; however, all pilidia tested by PCR for infection came from oocytes of a single *M. alaskensis* female. Only one visibly infected 2 - week-old pilidium was found positive by PCR. All other individuals tested negative by PCR and had no visible signs of infection.

The infected pilidium carried spores in mesenchyme cells, mobile cells which crawl and transport materials around the larval body. Other microsporidia are known to exploit mobile cell types for migration to specific tissues within the host. For example, in human patients, *Enterocytozoon bieneusi* infects macrophages, a cell similar in structure and function to larval mesenchyme cells. If intentional, infection of these migratory cells may permit transport into future juvenile tissues. Alternately, if indeed mesenchyme cells of the pilidium larva function like macrophages, these cells may be capable of voiding infection before spores have a chance to infect other larval tissues. An immunologic response may explain why infections were rarely found within developing pilidia and were never detected in newly metamorphosed juveniles.

#### 5.4.2 Horizontal transmission

In horizontal transmission, microsporidia are nearly always found inside the gut epithelia of infected hosts. The nemertean body plan includes gut diverticula which are interdigitated with gonads to promote diffusive nutrient transport. This anatomy lends itself to a strategy where spores maturing in the lumen of the gut could potentially puncture through gut and ovarian walls to directly infect an oocyte within an adjacent ovary. Infections in *M. alaskensis* seem to cluster around apparent epicenters and are not found in every ovary except in cases of heavy infection. High parasite loads observed in many specimens and the high prevalence found in Coos Bay *Maculaura* may be indicators of horizontal transmission. However, if the parasite is horizontally transmitted, spores should be found within the gut and, so far, we have found no evidence of microsporidia anywhere but within oocytes. Nevertheless, given the lack of evidence for successful vertical transmission, horizontal transmission to *M. alaskensis* is most likely, as it is the initial mode of transmission exhibited in microsporidian infections (Weiss and Becnel, 2014).

Based on the observation of chaetae within gut contents of dissected *Maculaura*, polychaete worms likely make up all or part of their natural diet (Maslakova and von Dassow, pers. obs.). *Armandia brevis* is a burrowing polychaete worm found in sandy tidal flats alongside *Maculaura* and is a potential prey for ribbon worms in these environments. In 1970, Szollosi discovered a microsporidian infecting the eggs of *A. brevis* in Friday Harbor, Washington. This microsporidian was classified as *Pleistophora sp*. based on morphological characters (Szollosi, 1970). It is diplokaryotic, develops within a parasitophorous vesicle, merogony proceeds by plasmotomy, and it has seven to eight turns of the polar tube in completed spores (Szollosi, 1970). Based on these morphological clues, the species found within *Maculaura* may be related to *Pleistophora* sp*. Pleistophora* sequences found in GenBank are not closely related to the Hepatosporidae but sequences of *Pleistophora* from eggs of *A. brevis* are not available for comparison. If it is closely related, then it may represent another member of the Hepatosporidae; if it is in fact the same species, it may be a source of infection for putative *Armandia* predators such as *Maculaura*.

### 5.5 Evidence of Novel Genus

Based on the discovery of unique SSU rRNA and RPB1 sequences, this microsporidian parasite likely represents not only a new species but also a new genus. The position on the SSU rDNA phylogeny suggests that it belongs to the family Hepatosporidae with *Hepatospora eriocheir* as the only other currently described member. A recent study by Tokarev et al. redefined the relationships between two micsorporidian genera. They found 91% sequence similarity among all members of the genus *Vaviomorpha* and 94% sequence similarity between members of the *Nosema*. Between the two genera, there was a 78% sequence similarity and a mean genetic distance of 0.183 (Tokarev et al., 2020). Direct comparison of SSU rDNA sequences *Oogranate pervascens* and *H. eriocheir* reveals between 76.61 and 76.71% sequence similarity and the genetic distance calculated by both the Bayesian and maximum-likelihood phylogenies is greater than 0.5.

When comparing morphological features between *Oogranate pervascens* and *H. eriocheir*, there are few shared characters (Table 3). The major difference is that *O. pervascens* produce very large PSVs which contain thousands of diplokaryotic spores. *H. eriocheir* produces small PSVs containing less than 60 unikaryotic spores (Stentiford et al., 2011; Wang and Chen, 2007). Based on comparisons of genetics and morphological characters these two species are not as closely related as one might expect between members of the same genus.

**Table 3.** Comparison of biological and morphological features of microsporidia in the family Hepatosporidae. Type habitat: F - freshwater host. M - marine host.

| Microsporidian species | Type host genus | Type habitat | Spores per PSV | Spore size and shape | Polar filament coils | Number of spore nuclei |
| --- | --- | --- | --- | --- | --- | --- |
| *Hepatospora eriocheir* | *Eriocheir* | Aquatic (F/M) | >60 | 1.8X0.9µm (ellipsoid) | 7-8 (single rank) | unikaryotic |
| *Oogranate* nov. gen. *pervascens* nov. sp. | *Maculaura* | Aquatic (M) | 10<sup>2</sup> – 10<sup>4</sup> | 2.3X1.3µm (ellipsoid) | 6-7 (single rank) | diplokaryotic |

### 5.7 Conclusion

We cannot declare with certainty how *Oogranate pervascens* infection of *Maculaura alaskensis* initiates nor can we thoroughly describe the relationship between the developing spores and the host cells. We can however hypothesize, based on our observations, that the evolutionary relationship between the microsporidian and its nemertean host may be proceeding towards a stable vertical transmission strategy in which the costs to the host are not substantial and do not cause overt signs of infection. Infection in other *Maculaura* species may look different than what was observed in *M. alaskensis*. Comparative studies of microsporidian infection among the other *Maculaura* species may provide insight into the evolutionary history of these congeners and of their microsporidian parasite which, at present, appears to be walking the line between horizontal and vertical transmission. It may offer clues for how and why this transition takes place and what it means for both host and pathogen.

## Acknowledgements

We would like to thank Dr. Maya Watts and Dr. Alan Shanks for their advice and their contributions to this research. Additionally, we would like to thank the professionals at the University of Oregon Histology Laboratory and CAMCOR facilities for their training and guidance.

## Notes

### Competing Interest Statement

The authors have declared no competing interest.

