## Supplementary figures and images for "A New Genus Of Microsporidian Parasite (Hepatosporidae; Micro-Sporidia) Found In The Oocytes Of Ribbon Worms From The North Pacific Genus Maculaura (Heteronemertea; Nemertea)"

### Supplemental File 1

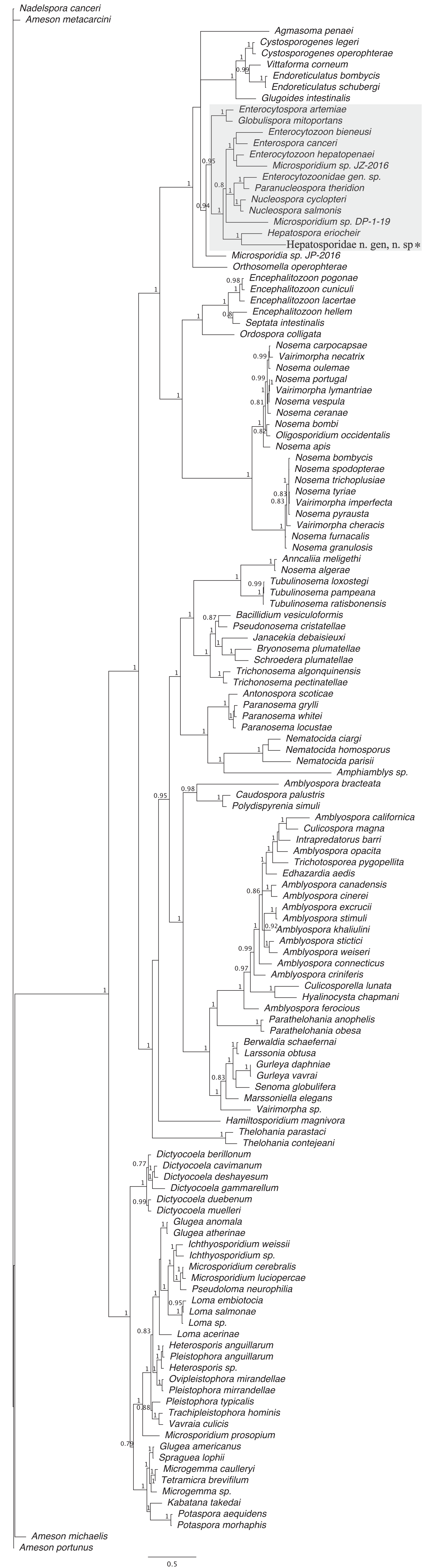

### Supplemental File 2

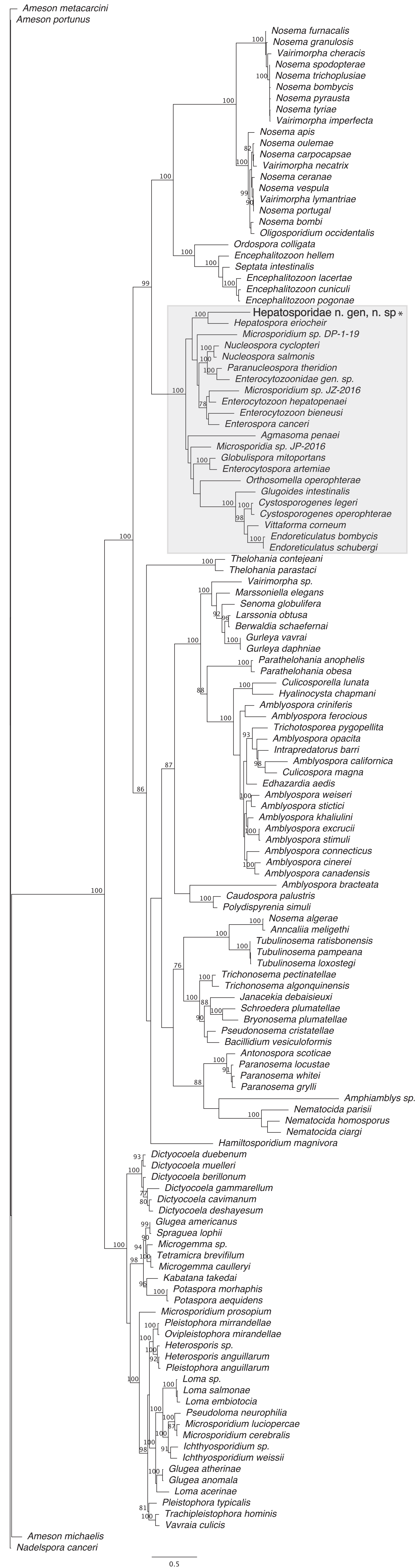
